# A Potential Role of Postsynaptic Limitation in Activity-Dependent Homeostasis

**DOI:** 10.1101/2025.10.20.683582

**Authors:** Vincent J. Chen, Peter G. Noakes, Nickolas A. Lavidis

## Abstract

The distribution of neurotransmitter release at amphibian neuromuscular junctions (**NMJ**) is characterised by a shift from Poisson to a tight binomial distribution as calcium concentration is increased. Despite advances in our understanding of neurotransmission, a paradox remains about how the large number of active zones (**AZs**) that contribute to release have such a low variance, resulting in a tight binomial distribution. Muscle-specific sodium channel-blocking neurotoxins have allowed us to examine the release characteristics when the unit size of transmitter release (quantal size) is not altered. Our present study has compared the neurotransmitter release characteristics when D-tubocurarine (curare) or the Na-channel blocking conotoxin (μ-conotoxin GIIIB) was used as the extracellular calcium concentration was increased. Quantal neurotransmitter release from toad *iliofibularis* motor nerve terminals was examined using intracellular electrodes and focal extracellular electrodes. Motor nerve terminal branches for recordings were located using DiOC_2_(5)-fluorescent imaging. Muscle action potentials were suppressed by using μ-conotoxin GIIIB (μ-CgTx GIIIB, 2.0 x 10^-6^ M) or curare, 6.5 x 10^-6^ M. Under μ-CgTx GIIIB, the maximum transmitter release reached was 25 quanta at high extracellular calcium concentration (2.1 mM); conversely, under curare, quantal content reached 148 quanta (*P*=0.0067). Using binomial analysis, it was found that this quantal increase was due to a significant increase in the number of release sites ‘n’ rather than the average probability of release ‘p’. Based on these observations, along with analyses of ECP shape and duration, we propose a mechanism in which a fast autoinhibitory response (< 5 ms from the nerve terminal impulse) limits the maximum level of depolarisation by voltage-dependent regulation of the acetylcholine receptor (**AChR**) channel conductance.

## Introduction

At the neuromuscular junction (**NMJ**), neurotransmitters are released as multiple packets (**quanta**) in a discrete step-like fashion. Tightly controlled quanta of acetylcholine (**ACh**) are released as an evoked response when the nerve terminal is depolarised by the arrival of an action potential or released spontaneously in the form of single quanta independent of presynaptic action potentials. Endplate potentials (**EPP**) are a measure of the evoked response composed of multiple quanta, while miniature endplate potentials (**MEPP**) are a measure of spontaneous single quanta releases (Del Castillo & Katz, 1954b; Fatt & Katz, 1952b). The exocytosis of neurotransmitters for both evoked and spontaneous responses occurs at electron-dense presynaptic release sites known as active zones (**AZs**) (Del Castillo & Engbaek, 1954; Del Castillo & Katz, 1954b). Ultrastructural studies reveal that hundreds of AZs are aligned perpendicular to the length of motor nerve terminal branches, opposite to the postsynaptic folds located on the membrane of the muscle fibres, where high densities of postsynaptic acetylcholine receptors (**AChRs**) are located at the lips of the postsynaptic folds (McMahan & Kuffler, 1971; Nishimune et al., 2004; Porter & Barnard, 1975). Within the nerve terminal, neurotransmitters are packaged into synaptic vesicles (**SVs**) by several proteins collectively termed vesicular-associated proteins. Many SVs are docked at each AZ, but only a single quantum of neurotransmitter is released per AZ in most cases (Del Castillo & Katz, 1954a; Meriney & Dittrich, 2013). Each quanta (single vesicle) can contain 2,000 molecules of acetylcholine (Hirsch, 2007). For amphibia, each NMJ has hundreds of AZs (Meriney & Dittrich, 2013), and each AZ contains double rows of 20 to 40 docked SVs (Heuser et al., 1974; Pawson et al., 1998). These docked SVs constitute the ‘ready releasable pool’ (**RRP**) (Shotton et al., 1979). With such a vast number of SVs the question arises, is the release of quanta tightly regulated to prevent overstimulation of the muscle?

Experiments conducted over the 1960s to 1980s found that extracellular calcium concentration (**[Ca^2+^]_o_**) is vital for evoked release, and that as [Ca^2+^]_o_ increases, the number of quanta released (quantal content) by a nerve impulse increases sigmoidally by a three to fourth power (Bennet & Lavidis, 1982; Bennett & Fisher, 1977; Bennett & Lavidis, 1979; Dodge & Rahamimoff, 1967; Ge et al., 2020; Jenkinson, 1957; Stevens, 2003). From these studies, it was predicted that as [Ca^2+^]_o_ increased, quantal content would increase following a sigmoidal relationship reaching an asymptote close to the total number of AZs (Bennet & Lavidis, 1982; Bennett & Fisher, 1977; Bennett & Lavidis, 1979; Dodge & Rahamimoff, 1967; Jenkinson, 1957). To prevent the initiation of the muscle action potential and subsequent contraction, the acetylcholine receptor competitive antagonist D-tubocurarine (**curare**) was used to reduce the quantal size and thus prevent the EPP from reaching the threshold for initiating the muscle action potential. We asked the question, what would be the shape of the sigmoidal relationship between [Ca^2+^]_o_ and quantal content if the quantal size was not reduced with curare? With the introduction of the muscle-specific Na-channel blocking conotoxin, it became possible to examine the sigmoidal relationship between [Ca^2+^]_o_ and quantal content without the need to reduce quantal size.

The relationship between quantal size and quantal content has gained interest recently, with the observation that curare, vecuronium, and alpha-bungarotoxin (**α-BTX**) were all able to significantly increase quantal content (Wang et al., 2018). Furthermore, Wang and colleagues found that the increase in quantal content could be attributable to an increase in the number of SVs released from AZs. From these observations, it was concluded that AChRs may have the ability to signal to the presynapse via noncanonical signalling, thereby preventing the release of additional SVs. Here, an alternative hypothesis involving the characteristics of the AChRs is proposed to explain possible autoregulation of the postsynaptic depolarisation. A postsynaptic contribution is likely, considering the decrease in endplate current half-life observed in their study (Wang et al., 2018). Curare’s effect of reducing the size of the quanta could allow more evoked quanta to be observed; these would be quanta that are normally released and go unnoticed by the level of depolarisation (the idea of missing quanta) (Nguyen & Stewart, 2016; Wojtowicz et al., 1994). Previous estimates of quantal content from EPP amplitude measurements as extracellular calcium concentration is increased may not reflect the actual number of evoked quanta released (Bennet & Lavidis, 1982; Bennett & Fisher, 1977; Bennett & Lavidis, 1979; Dodge & Rahamimoff, 1967; Jenkinson, 1957). It is possible that postsynaptic excitability is autoregulated by the characteristics of the AChR related to its depolarisation-induced inactivation (Gage, 1976; Sine & Steinbach, 1984; Vyskocil et al., 2009). Thus, by reducing quantal size with curare, more evoked quanta are observable.

Another mechanism that may regulate postsynaptic excitability is via presynaptic autoinhibition involving the neurotransmitters released. The discovery that neurotransmitter release is non-uniform (Bennet & Lavidis, 1982; Bennett et al., 1986; Bennett & Lavidis, 1979; D’Alonzo & Grinnell, 1985; Dengyun Ge, 2018; Dengyun Ge et al., 2020) supports the idea that highly active release sites may autoinhibit release from adjacent active zones and thus contribute to non-uniformity in neurotransmitter release (Ge et al., 2018; 2020). The possibility that a fast autoinhibitory system may exist along nerve terminal branches was proposed by Bennett and Robinson in 1990 and was further expanded by Ge et al in 2020 into a conceptual model, wherein the relative activity of AZs could be established by the most active release sites (AZ’s) imposing a surrounding autoinhibition. Thus, while a nerve terminal may be composed of ∼ 700 AZs, it behaves as 25 release units, with each release unit composed of 40 AZs acting together. In previous interpretations of this proposed model, it had been assumed that the local inhibitory mechanism occurs via presynaptic autoregulation involving transmitter-induced negative feedback (Ge et al., 2020). Given the contemporary focus upon neurotransmission as not simply synaptic release but as a tightly associated trans-synaptic complex, we propose that local autoinhibition could contribute to limitation by postsynaptic excitability.

To test this idea, the present study employed the use of µ-conotoxin GIIIB (**µ-CgTx GIIIB**), a muscle-specific voltage-gated Na_v_1.4 channel blocker, which blocks muscle contractions without altering quantal size (Magown et al., 2015b), but does not affect the propagation of the action potentials to the nerve (i.e., no blocking effects of axonal sodium channels; (Scharman, 2005)). These features of µ-CgTx GIIIB allowed us to compare the relationship between quantal content and extracellular calcium concentration of amphibian NMJs in the presence of curare and µ-CgTx GIIIB.

## Methods

### Tissue preparation

Adult cane toads, 45-55 mm in length from snout to ischium, were collected from Southeast Queensland, Australia. Toads were maintained overnight in a humidified tank with a 12/12-hour light-dark cycle at seasonal outdoor temperatures. Animals were culled through double pithing according to the Animal Ethics Committee of the University of Queensland (2023/AE000468).

The *iliofibularis* muscle and innervating nerve were dissected free from the surrounding connective tissue, tendinous insertions, and sciatic nerve. The muscle was pinned onto a bed of cured silicone rubber (Sylgard, Dow Corning Corp., MI, USA) within a 4ml capacity organ bath. The preparation was then transferred to the recording stage of an Olympus BH2 upright microscope, where the bath was perfused with room temperature Ringer solution with the following composition (mM): Na^+^, 147.83; K^+^, 4.7; Mg^2+^, 1.2; Cl^-^, 130.8 – 134.5; H_2_PO_4_^-^, 0.67; HCO_3_^-^, 23.96; Ca^2+^, 0.25 – 2.1; glucose, 7.8. O_2_ (95%) at a rate of 3-5 ml per minute. CO_2_ was continually dissolved in the solution through bubbling at 18 ± 2 °C, and maintained a pH between 7.3 and 7.4. [Ca^2+^] (0.25mM to 2.1mM) and [Mg^2+^] (1.2mM) were supplied through Ca_2_Cl and Mg_2_Cl dissolved in the Ringer’s solution supplying the bath. All chemicals were sourced from Sigma-Aldrich (St. Louis, MO, USA). Stimulation at 0.17 Hz was delivered for 30 - 40 minutes to wash out endogenous calcium and equilibrate the Ringer solution [Ca^2+^]_o_.

The *iliofibularis* nerve was drawn into a glass pipette using gentle negative pressure. The pipette acted as the stimulating electrode through two silver chloride wires, one within the pipette as a positive electrode, and the other wrapped outside as the negative electrode. A current pulse of 0.1 ms (duration) and 1-10 V (amplitude) at 0.2 Hz (frequency) was delivered through the electrode to stimulate the nerve-muscle preparation using a Grass Instruments stimulator (SD9) coupled to a Grass stimulus isolator (SIU5).

### Intra- and extra-cellular electrophysiological recordings

Once an appropriate NMJ site was located to within 0.5 mm from the intracellular recording electrode, stimulation of the muscle nerve was turned off for 5 minutes to allow for vesicular replenishment and avoid stimulus-induced conditioning. A stimulus train of 200 evoked end-plate potentials was recorded at each impaled site, with at least 30 spontaneous end-plate potentials sampled. The rise time of the recording served as an indicator for electrode proximity to the nerve terminal or nerve terminal branch in the case of extracellular recordings. Intracellular recordings were aborted if: the initial resting membrane potential of the impaired skeletal muscle cell fluctuated by more than 10% during the recording period, the frequency of spontaneous end plate potentials increased by more than 15%, or if the skeletal muscle fibres were twitching. Extracellular recording was accepted for analysis if: the nerve terminal impulse (**NTI**) was clearly visible; the rise time of the end plate current was less than 2 ms; and the frequency of the spontaneous end plate currents did not vary by more than 15% (Dengyun Ge, 2018). These recordings were amplified using an Axoclamp 2B amplifier (Molecular Systems, Sunnyvale, CA, USA) and digitised at a 10kHz sampling rate using the MacLab system and Scope software (Version 3.5.5, AD Instruments, CO, USA).

For intracellular electrophysiology recordings, sharp borosilicate glass microcapillary electrodes filled with 2M KCL (50 – 80 MΩ) were used as per our previous studies (Bennett & Lavidis, 1979). A rise time of less than 2ms for evoked and spontaneous/miniature end plate potentials (**EPPs** or **mEPPs**), along with a resting membrane potential (**RMP**) between -60 to -80mV, was considered to be an acceptable recording. RMP was monitored continuously. For extracellular recordings, glass microelectrodes with a tip diameter of 3-5 μm filled with 0.5M [Na^+^] Ringer solution was used.

To locate and map the motor nerve terminals, the *iliofibularis* nerve-muscle preparation was bathed for 1-2 minutes in 0.1μM of 3,3’-diethyloxardicarbocyanine iodide (**DiOC_2_(5)**) in the dark, and then washed for 3 minutes with Ringer’s solution, as per our previous studies (Bennett et al., 1986; Dengyun. Ge & Lavidis, 2017; Lavidis et al., 2008). These stained nerve terminals were fluoresced at 555nm using an Olympus BH2 microscope and image intensifier camera (Panasonic) and displayed on a video monitor. The outline of the fluorescent terminals was traced onto the monitor screen. Fluorescence was then switched off, and structural reference markers, including the outline of the viewed nerve terminal branch, were traced onto the screen. This process allowed us to place an extracellular electrode at discrete sites along the length of the imaged terminal that were separated from neighbouring branches. The microelectrode was lowered to the muscle membrane containing the motor nerve terminal, to form a focal loose patch enabling us to record the NTI, evoked and spontaneous/miniature endplate currents (**EPCs** and **mEPCs**), respectively. Miniature/spontaneous (**mEPC**) frequency was monitored to control for stimulation emanating from electrode pressure (Fatt & Katz, 1952a).

### Calcium dependence

The extracellular calcium concentration ([Ca^2+^]_o_) was altered by changing the concentration of calcium chloride dissolved in the reservoir of the Ringer’s solution supplying the nerve-muscle preparation. The [Ca^2+^]_o_ used in this investigation ranged from 0.25mM to 2.1mM. The preparation was first incubated for 30-40 minutes in the lowest concentration used to remove endogenous levels of calcium present in the muscle. Extracellular calcium concentration was then increased, with magnesium concentration at a constant 1.2 mM to prevent possible changes in the conduction of nerve impulses due to divalent cation concentration falling below 0.7 mM (Frankenhaeuser & Hodgkin, 1957).

### Toxin preparation and application

Synthetic polypeptides µ-conotoxin GIIIB (**µ-CgTx GIIIB**) were purchased from Alomone Laboratories (Jerusalem, Israel). D-tubocurarine Chloride (**Curare**) was purchased from Sigma-Aldrich (St. Louis, MO, USA). 6.5 x 10^-6^ M curare or 2 x 10^-6^ M of µ-CgTx GIIIB was added to the reservoir perfusing the organ bath (Bennett et al., 1977; Dodge & Rahamimoff, 1967; Li & Tomaselli, 2004). The preparation was left in the modified solution for forty minutes or until no more contraction could be observed (averaging 30 minutes). Carbogen (5% carbon dioxide, 95% oxygen) was perfused directly into the bath.

### Quantal content calculations

EPPs and MEPPs with amplitude greater than twice the height of the average noise level were identified as evoked or spontaneous signals. Intracellular quantal content (***m_i_***) was calculated using the formula (Bennett and Lavidis, 1979): 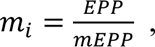 where EPP is the average amplitude of the end-plate potential and mEPP is the average amplitude of the miniature end-plate potential. When failure to evoke quantal release accounted for more than 30% of cases, we employed the method of failures (Boyd & Martin, 1956): 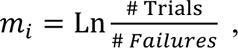 where intracellular quantal content is equal to the natural log of the number of stimulations (evoked and failures) divided by the number of failures (Boyd & Martin, 1956). Extracellular quantal content (***m_e_***) was calculated by the total number of quanta released (singles, doubles, triples) divided by the total number of trials (stimulations) (Ge & Lavidis, 2017): 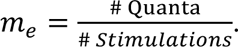

### Binomial parameter calculations

Binomial parameter “p” was determined using methods from Bennett and Lavidis 1979, where: 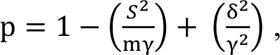 where m and S^2^ are the mean and variance of EPP amplitudes, and γ and δ^2^ are the mean and variance of mEPPs. Meanwhile, the binomial parameter “n” was determined from: 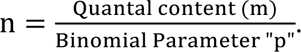 Curarised preparations require correction for accurate estimation of amplitude and *m_i_* according to the formula:

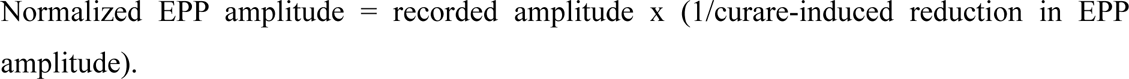

### Decay half-life calculations

For extracellular potentials (EPCs and mEPCs), decay time (**t½**) was determined by fitting an exponential decay curve according to the exponential decay formula (Magleby and Stevens, 1972): *y*(*t*) = *ae*^*kt*^, where *y*(*t*) is the value at time “*t*”, a is the amplitude of the EPC, “*k*” is the rate of decline, and “t” is time. Half-life was then calculated using the formula: 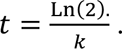

### Calcium dependence calculations

Calcium dependence was determined by plotting quantal content on double logarithmic coordinates, where the resulting slope represents the power relationship. Quantal content and [Ca^+2^]_o_ were also plotted on a double-reciprocal coordinate to form a Lineweaver-Burk plot (Dodge & Rahamimoff, 1967). Linear regression was then applied to the data points. This plot allowed for the estimation of the peak evoked response possible at the NMJ, at a theoretical infinite [Ca^2+^]_o_ by looking at the y-intercept. To correct quantal content for non-linear summation of units, equations from Martin (1955) and MacLachlan and Martin (1981) were used (Martin, 1955; McLachlan & Martin, 1981). These equations were used to analyse data from Bennett et al (1977) and Dodge and Rahamimoff (1967) to a maximum quantal content meta-analysis (**Table 1**).

**Table 1.**
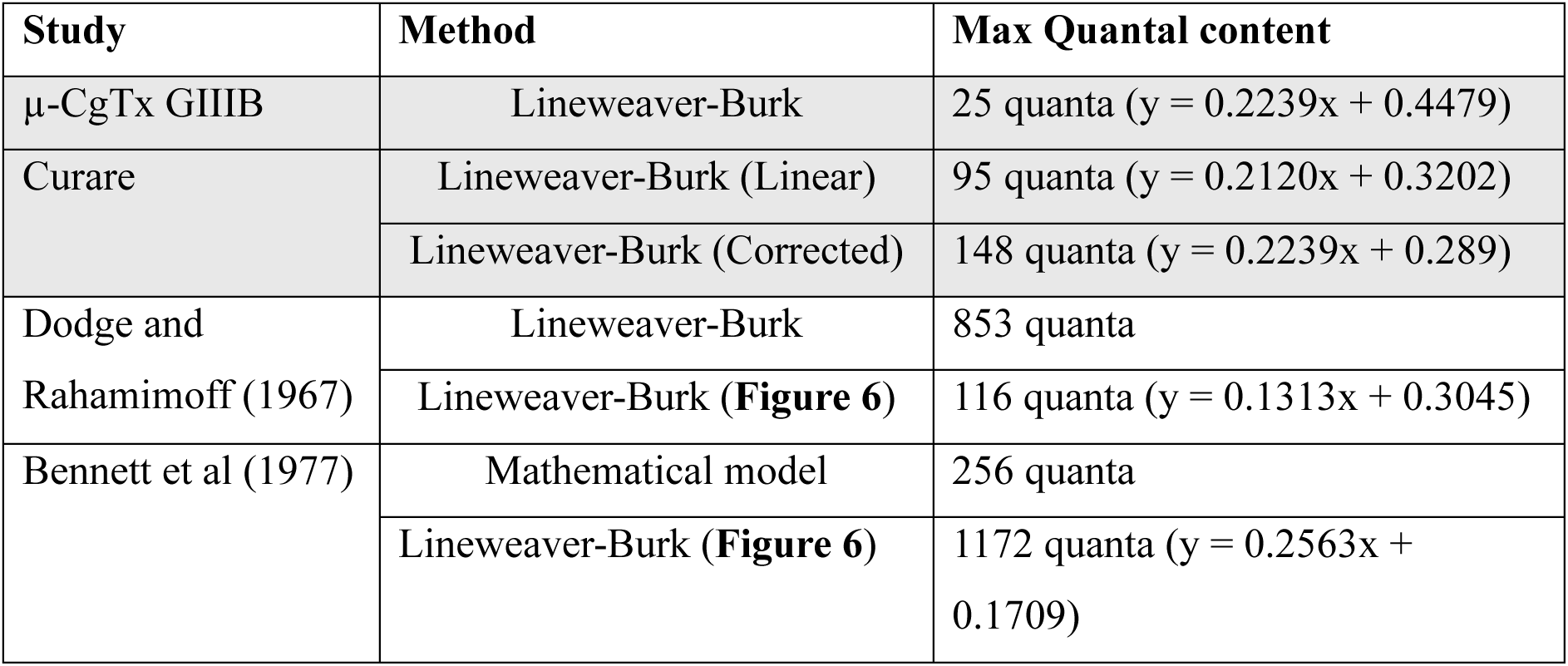
Table of maximal quantal release found in the literature. Maximum quantal content was determined using graphs from literature, and analysis by past researchers (methods). To compare with results from the present study, a Lineweaver-Burk analysis was performed on experimental data sets most similar to our experimental conditions, from these studies. Cells shaded grey are results from the present study.

A sigmoid curve (see **Figure 2A**) was generated from our experimental data using the Dodge-Rahamimoff formula (cited above (Dodge & Rahamimoff, 1967)). This data was extrapolated to a high [Ca^2+^]_o_ of 10mM using the Michaelis-Menten formula at a fourth power as follows: 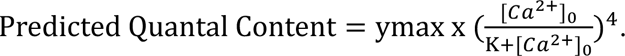 Where “ymax” is the y-intercept (maximal quantal content) derived from the double reciprocal plot, and “K” is the dissociation constant derived from the x-intercept.

### Significance tests

Two-way ANOVA with Tukey’s HSD test was used to conservatively control for family-wise error across multiple groups. Given heterogeneity in sample sizes and across included studies (Figure 6), a conservative test was preferred to minimize false positives. Student’s *t*-test were used to determine significance. "Post-hoc pairwise comparisons " Data are expressed as mean ± standard error of the mean (**SEM**) with statistical significance accepted at *P* < 0.05. LabChart and Scope S060/7 (AD Instruments) were used for data analysis.

## Results

### The sigmoidal relationship between end plate potential amplitude and [Ca^2+^]_o_ shows a lower asymptote limitation of release under µ-CgTx GIIIB compared to curare

To determine whether there was a limitation of transmitter release, we characterised the effects of curare and µ-CgTx GIIIB. The relationship between EPP and [Ca^2+^]_o_ was examined under these two toxins, using intracellular recordings while simultaneously recording MEPPs. As we increased [Ca^2+^]_o_ to beyond 0.4 mM, EPPs and MEPPs were recorded using µ-CgTx GIIIB to prevent the generation of muscle action potentials and muscle contractions (Magown et al., 2015a). Superimposed, consecutive EPPs and MEPPs are shown in **Figure 1A** for [Ca^2+^]_o_ of 0.4mM (left) and 0.9mM (right). EPPs were identified following the stimulus artefact at both concentrations, where the amplitude of the EPPs increased when the [Ca^2+^]_o_ was raised, from 0.4mM to 0.9mM. Failures were observed in 0.4 mM [Ca^2+^]_o_, but became absent as [Ca^2+^]_o_ was increased. Examples of MEPPs and EPPS are shown in **Figure 1A** (i.e., MEPPs = grey traces; EPPs = black traces). MEPP amplitude remained consistent at 0.84 +/- 0.025 mV irrespective of [Ca^2+^]_o_.

**Figure 1.**
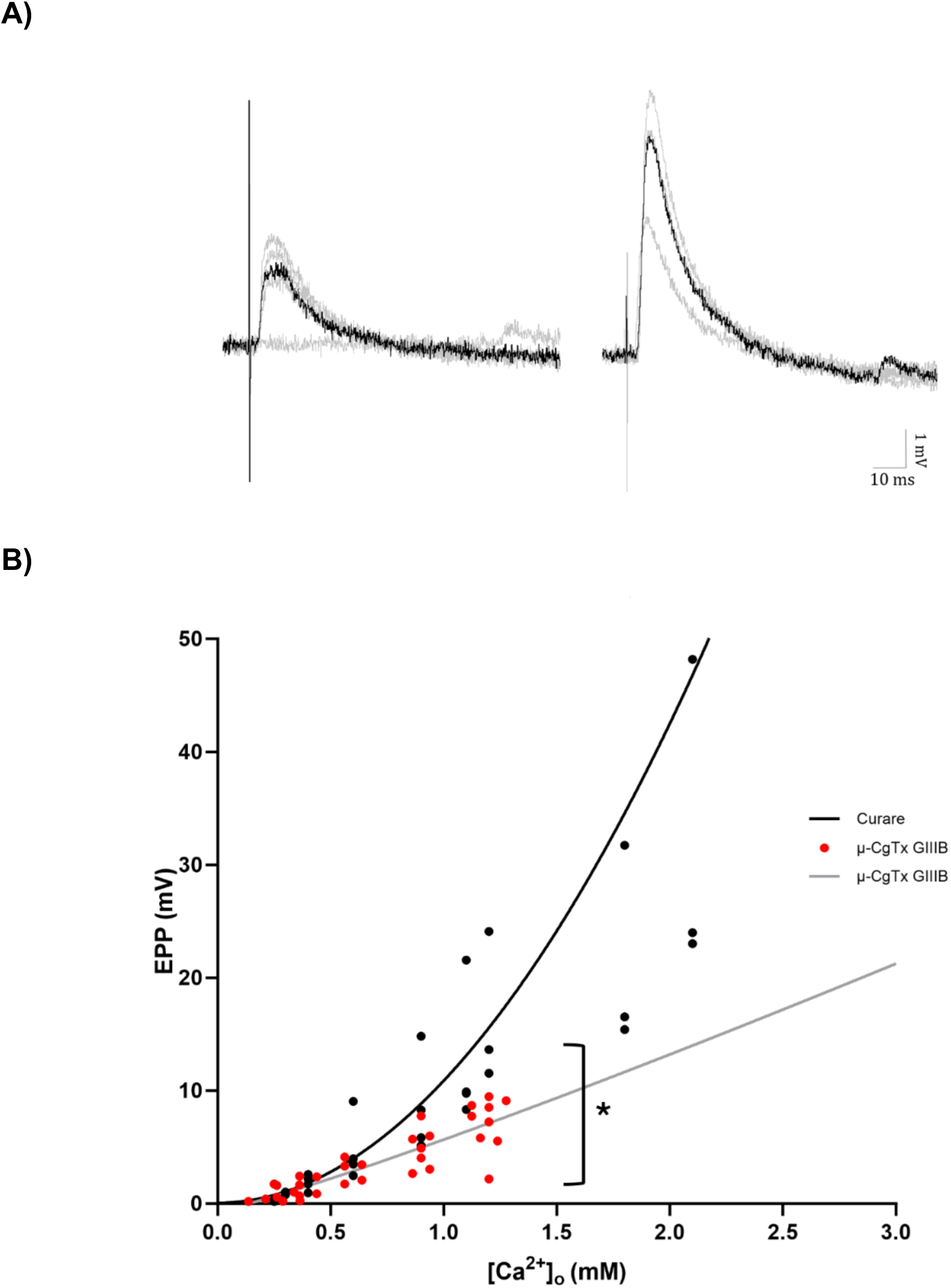
The relationship between [Ca2+]_o_ and evoked EPP amplitude shows a disproportionate increase under curare compared to µ-CgTx GIIIB. **A)** Examples of intracellular recordings of EPPs and MEPPs shown overlayed consecutively. Nerve-muscle preparations perfused with 0.4mM (*left*) or 0.9mM (*right*) [Ca^2+^]_o_ treated with µ-CgTx GIIIB (2 μM). B) Relationship between [Ca^2+^]_o_ and EPP amplitude for µ-CgTx GIIIB (*n*=12) and curare treatments (*n*=5). Nerve-muscle preparations perfused with 0.25 to 2.1mM [Ca^2+^]_o_ in the presence of curare (black data points) and 1.2mM µ-CgTx GIIIB (red data points). Despite a smaller quantal size (MEPP amplitude), the application of curare resulted in a significantly higher EPP amplitude as [Ca^2+^]_o_ increased (*P* =0.0246, *, two-way ANOVA). The black and grey trendlines represent fitted sigmoidal dose-response curves.

The control study at 0.25mM to 0.4mM of [Ca^2+^]_o_ demonstrated no significant difference in the EPP amplitudes across the two toxin treatments (i.e., curare Vs µ-CgTx GIIIB; *P*=0.75; **Figure 1B**). However, as [Ca^2+^]_o_ increased, this difference became more divergent (*P*=0.021). For µ-CgTx GIIIB treatment, there was an increase from 1.61± 0.37mV at 0.4mM [Ca^2+^]_o_ to 7.15 ± 0.77 mV at 1.2 mM [Ca^2+^]_o_, representing a <u>444%</u> increase. By contrast, for curare treatment there was an increase from 1.88 ± 0.35mV at 0.4mM [Ca^2+^]_o_ to 16.43 ± 3.88mV at 1.2mM [Ca^2+^]_o_, which represented an <u>875%</u> increase (**Figure 1B**). Taking our experimental results to plot a sigmoidal dose-response curve revealed that muscles treated with curare had a steeper slope (1.977, R^2^=0.72) compared to µ-CgTx GIIIB (1.137, R^2^=0.75) (**Figure 2A)**.

**Figure 2.**
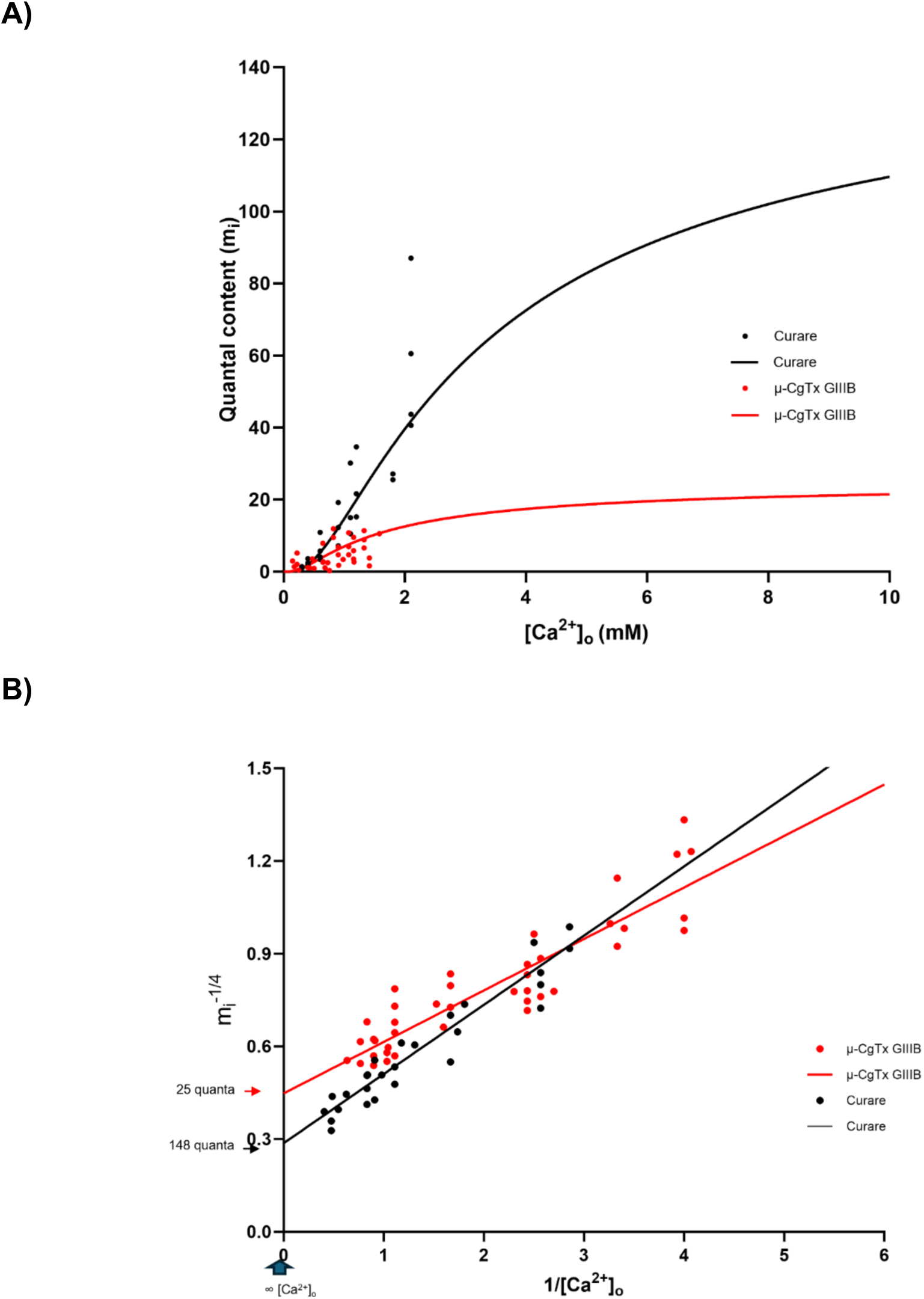
Relationship between [Ca2+]_o_ and quantal content (m*_i_*) shows a higher maximum possible quantal content under curare compared to µ-CgTx GIIIB. **A)** Quantal content under curare and µ-CgTx GIIIB plotted on double log coordinates. A sigmoidal curve is fitted to the data using a fourth power relationship from Dodge and Rahamimoff (1967), see methods (*P* = 0.0067, **, two-way ANOVA). **B)** Double reciprocal relationship between [Ca^2+]^_o_ and quantal content (m*_i_*) for µ-CgTx GIIIB (*n*=12) and curare treatments (*n*=5). At an infinite concentration of [Ca^2+]^_o_, µ-CgTx GIIIB m*_i_* was limited to 25 quanta content, while the curarised amplitude was 148 quanta (*P* = 0.0023, two-way ANOVA). Nerve-muscle preparations perfused with 0.25 to 1.2mM [Ca^2+]^o treatment with curare (black) and µ-CgTx GIIIB (red).

Altogether, these findings revealed that the application of curare resulted in a significantly higher EPP amplitude when compared to µ-CgTx GIIIB across all concentrations tested (*P*=0.0226).

### The sigmoidal relationship between quantal content and [Ca^2+^]_o_ shows a lower asymptote limitation of quanta content under µ-CgTx GIIIB compared to curare

To better estimate the maximum possible neurotransmitter release under curare and µ-CgTx GIIIB, we used mathematical models from Dodge and Rahamimoff 1967 (see methods), and a double reciprocal plot. When a trend was extrapolated based on fourth power mathematical models (Dodge and Rhamimoff 1967), it was revealed that in the relationship between quantal content and [Ca^2+^]_o_, a lower asymptote was reached when µ-CgTx GIIIB was used. While bathing the muscle in curare resulted in a sigmoidal curve for quantal content versus [Ca^2+^]_o_, it indicated a significantly higher asymptote, by contrast. This result indicated an inverse relationship between quantal size and quantal content (**Figure 2**).

While quantal content increased as [Ca^2+^]_o_ increased in both treatments, quantal content under µ-CgTx GIIIB increased at a slower rate, compared to curarised treatment (*P* = 0.0067, **Figure 2A**). A double reciprocal plot of quantal content (mi^-1/4^) versus [Ca^2+^]_o_ (**Figure 2B**) estimated the quantal content at the y-intercept (when [Ca^2+^]_o_ approaches infinite concentration) for curarised preparations at mi^-1/4^ = 0.2868 (148 quantal content) and for µ-CgTx GIIIB, mi^-1/4^ = 0.4479 (25 quantal content, *P* =0.0023, **Figure 2B**). This may indicate that curare causes neurotransmission to become upregulated or less limited.

### Application of curare increases the number of release sites that participate in quantal release

To attempt to identify the mechanism behind this upregulation (or removal of inhibition), a binomial analysis was conducted using the EPP and MEPP amplitudes versus the number of observations to determine the number of active release sites (**n**) and the average probability of quantal release (**p**) at each recording site.

As can be seen in Figure 3, both the average probability and number of release sites increased as calcium concentration increased (**Figure 3A** and **3B**, respectively). However, for both µ-CgTx GIIIB (*P* = 0.95 ± 0.03) and curarised (*P* = 0.89 ± 0.09) preparations, the probability of release rapidly increased close to unity (i.e., p close to 1) by 0.7mM [Ca^2+^]_o_ (*P* = 0.6397, two-way ANOVA; **Figure 3A**). Consequently, any further increase in quantal content in concentrations higher than 0.7mM [Ca^2+^]_o_ as seen in **Figure 3C** was likely to be due to an increase in the number of participating release sites, as supported by the data in Figure 3C. Namely, the number of release sites (**n**) continued to increase from 7.78 ± 1.92 at 0.7mM [Ca^2+^]_o_ to 23.84 ± 5.71 at 1.3mM [Ca^2+^]_o_ under curare treatment; and an increase from 7.00 ± 1.04 at 0.7mM [Ca^2+^]_o_ to 13.40 ± 2.98 at 1.3mM [Ca^2+^]_o_ under µ-CgTx GIIIB treatment (**Figure 3B**). These changes represent an increase in the number of release sites by <u>306.4%</u> under curare, and <u>191.4%</u> under µ-CgTx GIIIB (*P*=0.0514, two-way ANOVA).

**Figure 3.**
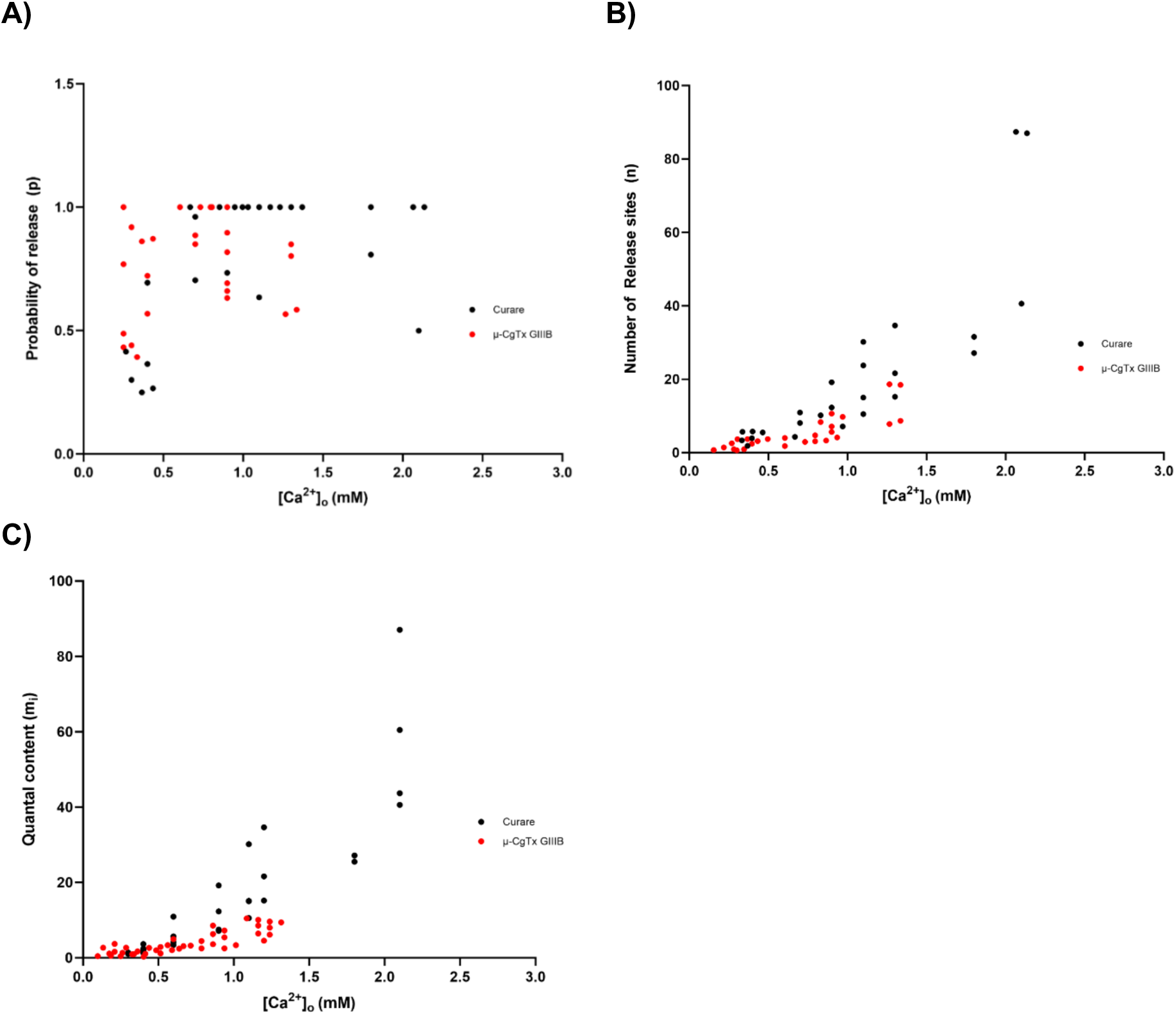
Binomial analysis of evoked release from terminals exposed to curare (n=4) or µ-CgTx GIIIB (n=7). The relationship between [Ca^2+^]_o_ and evoked quantal transmitter release. The decrease of quantal size (MEPP amplitude) by curare increased the number of evoked quanta released as [Ca^2+^]_o_ increased. **A)** The average probability of quantal release (p) showed an initial increase but reached unity by 0.7mM [Ca^2+^]_o_ while **B)** the increase in quantal content observed with curare was primarily due to an increase in the number of active release sites participating in release (n). Resultant from p and n, **C)** in the presence of curare, more quanta were released compared to the Na^+^-channel blocker to achieve the desired depolarisation for action potential activation (*P* = 0.0067, two-way ANOVA).

Thus, the increase in quantal content by either treatment can be explained predominantly by an increase in the number of a few active release sites reaching a probability of quantal release close to unity; a conclusion that is consistent with non-uniformity of quantal transmitter release (Bennett & Lavidis, 1979).

### Possible role of the acetylcholine receptor characteristics in controlling the duration and amplitude of the end-plate currents

Our observations using μ-CgTx GIIIB demonstrate that, in the absence of curare, there exists a certain kind of mechanism that limits the amount of evoked transmitter release to a maximum of 25 quanta. While the nature of this mechanism is not understood, literature suggests that the binding of curare induces a postsynaptic conformational change in acetylcholine receptors that alters the dynamics of endplate currents (i.e., the amplitudes of end-plate currents (Wang et al. 2018). It is therefore possible that such a postsynaptic mechanism (conformational change) could limit quantal release. Our data described below supports this idea:

When an extracellular electrode was placed within 1 to 2 μm from the nerve terminal, the electrode recorded a negative potential change, which had the same time course as patch electrode records when ACh is applied to the patch membrane containing acetylcholine receptors (**AChRs**). This signal, recorded by the extracellular electrode, is called the end-plate current (**EPC**), and detects the flow of Na^+^ ions into muscle through opened AChRs. Whereas the shape of the EPC reflects the stochastic nature of the AChR opening and closing times (Colquhoun & Sakmann, 1981; Kandell et al., 2021). When we increased [Ca^2+^]_o_, multiple quantal EPCs (as seen in **Figure 4A**) were recorded in proximity within the latency period for quantal release following the arrival of a nerve terminal impulse (Augustine & Kasai, 2007). We have observed that the amplitude of subsequent quanta following the first quanta released is often smaller in amplitude and has a shorter duration. We measured decay time (**t½**) changes between twin releases of EPC to determine if briefer conductance may be a factor in the limiting of postsynaptic depolarisation (**Figure 4A**).

**Figure 4.**
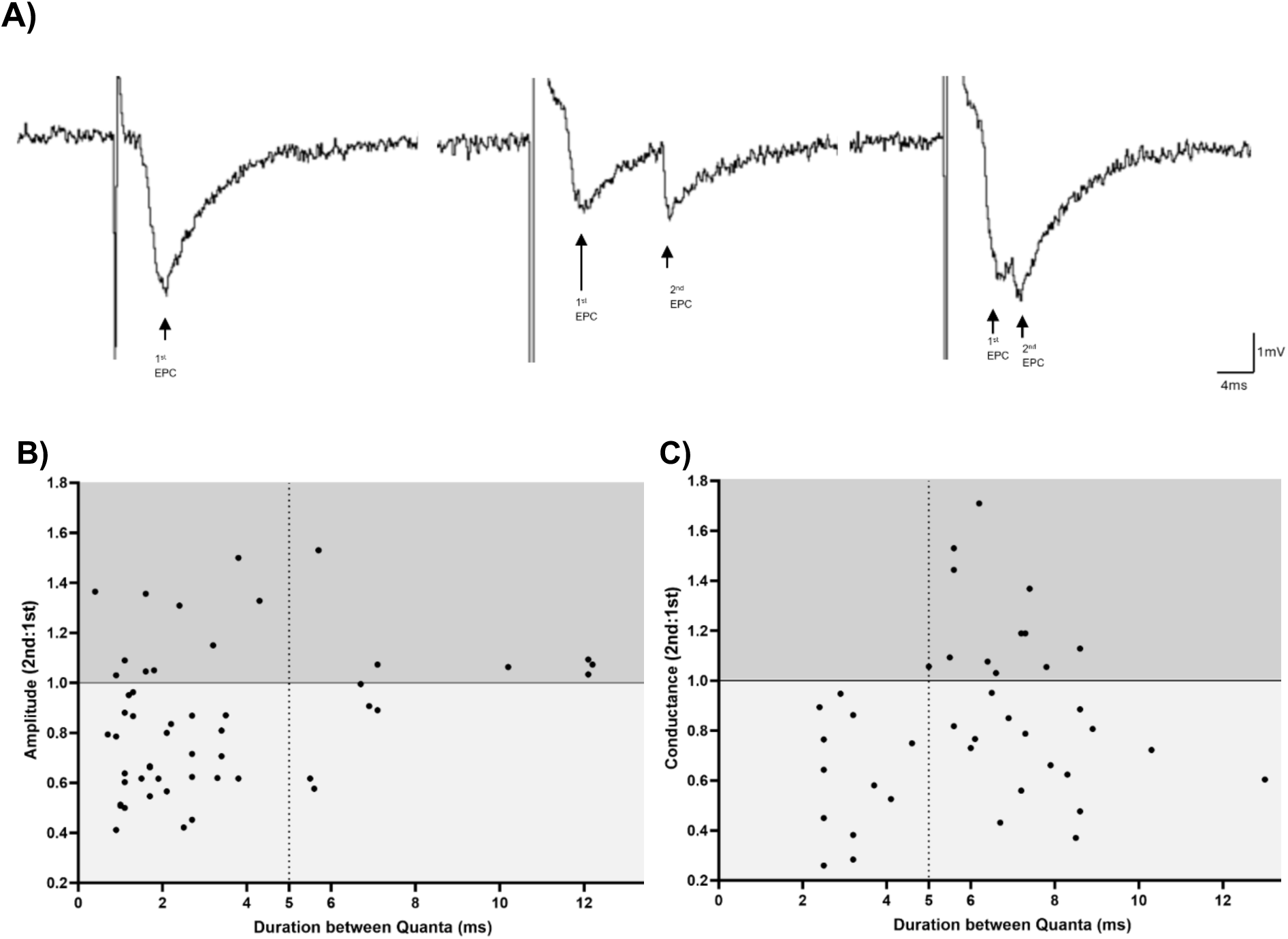
Autoregulation of evoked potential conductance at highly active release sites (n=10). **A)** *Left*: Example recordings of an EPC from an extracellular site where the electrode was placed approximately 2 mm to the side of a terminal branch visualised using DiOC_2_(5)-fluorescence. *Middle*: Example of a preceding current with an interval of 3ms between a first and a second EPC. *Right*: Example of a preceding current with an interval of 7ms between a first and a second EPC. Nerve-muscle preparations perfused with 0.4mM [Ca^2+^]_o_. **B)** The first end-plate current alters the ratio of time course and **C)** the ratio of amplitude of a second end-plate current if the interval between them is less than 5ms. The rate of recovery following the peak of the end-plate current was significantly shorter in the second end-plate current when compared with the recovery from the first end-plate current. This suggests that the AChRs may have a shorter open time in the second end-plate current and may be influenced by the previous opening in the first end-plate current.

By taking the ratio of the amplitude or decay of the first and the second EPC, any value below 1.0 would be an indication that the second EPC had a lower amplitude and a briefer conductance. We found that for all measured EPCs, if the duration between the first and the second EPC was less than 5ms, then conductance was lower in the second compared to the first (all points with a ratio of less than 1.0) **(Figure 4B)**.

A similar observation can be made for amplitude, where when paired EPCs were within 5ms of one another, the average amplitude ratio was below 1 (i.e., the second EPC smaller than the first) at 0.81±0.05. Whereas, when paired EPCs were further apart (>5ms), the amplitude ratio increased to 1.04±0.05 (i.e., the second EPC was identical to the first). This constituted a significant positive trend (*P*=0.006, *t*-test) **(Figure 4C)**.

Additionally, we observed a phenomenon at the toad NMJ wherein trains of multiple EPCs and mEPCs occur one after another in a “sawtooth” like formation - representative sample profiles of this occurrence can be observed in **Figure 5A**. We found that the first EPC is usually larger than the following EPCs. As shown in **Figure 5B**, the second EPC in a chain was smaller than the first by 27.30±5.01%, the third by 26.18±4.86%, the fourth by -38.50±4.88%, the fifth by 18.00±12.12%, and the sixth by 21.84±0.13%. This constitutes a significantly (*P*=0.0003) negative trend (y = -0.06602x + 0.9612).

**Figure 5.**
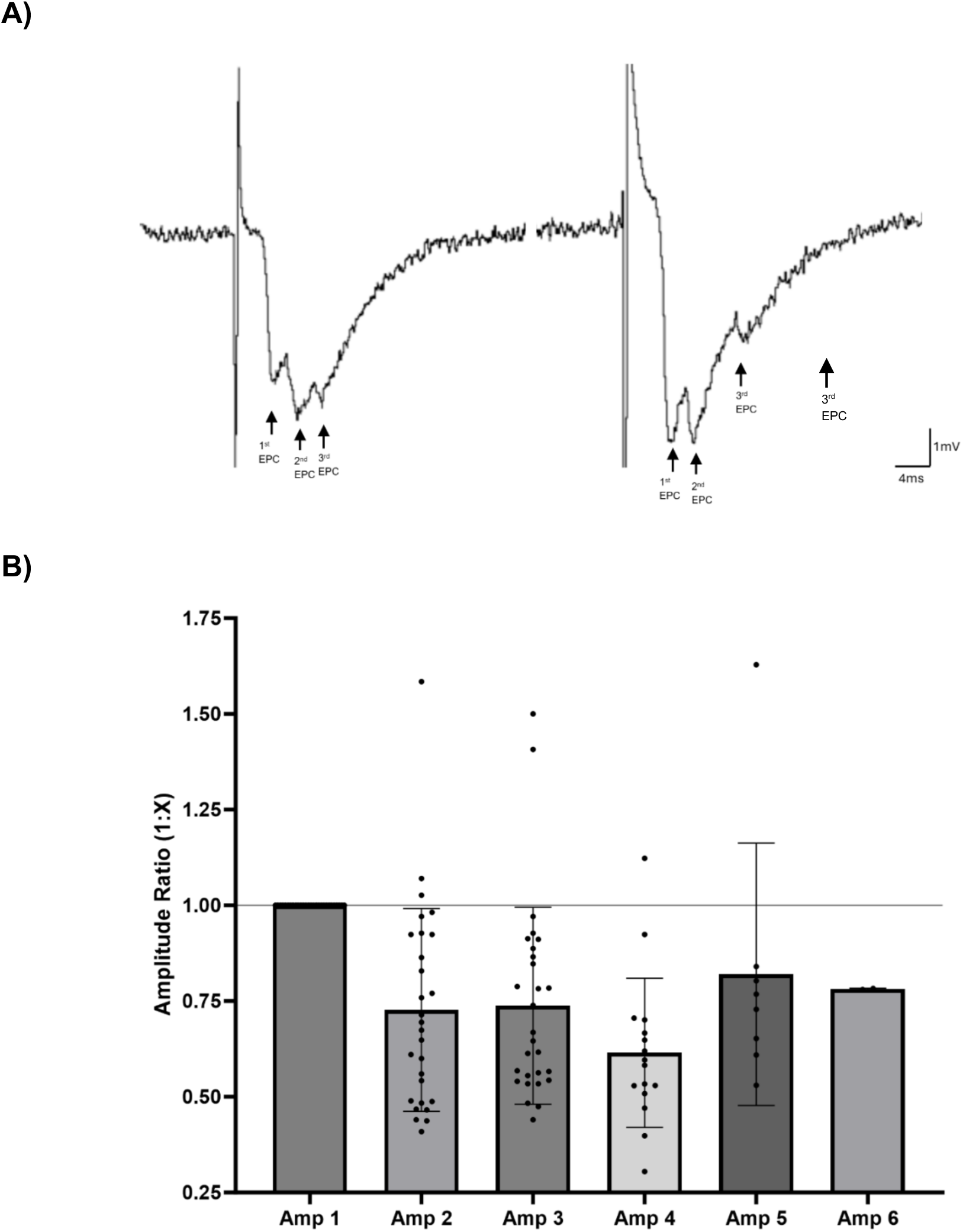
The amplitude of multiple end plate current release events (n=6). **A)** Example recordings of EPCs by an extracellular recording electrode placed approximately 2 mm to the side of a DiOC_2_(5)-fluorescent visualized terminal branch. *Left EPC trace*: Example of a preceding current with an interval of 3ms between a first and a second EPC. *Right EPC trace*: Example of a preceding current with an interval of 7ms between a first and a second EPC. Nerve-muscle preparations perfused with 0.4mM [Ca^2+^]_o_. **B)** The amplitude of subsequent end-plate currents 2 to 4 was significantly reduced when compared to the amplitude of the first end-plate current (*P*<0.005, t-test). This constituted a significant negative slope (P=0.0003) (y = -0.06602x + 0.9612). Multiple quantal releases were collected for analysis. Each train had to be separated from any previous quantal release by at least 1 second. Correction for non-linear summation was applied. Error bars indicate SEM.

Together, these findings raise the possibility of latency and thus receptor characteristics playing a role in naturally limiting excitability. This will be further discussed below.

## Discussion

This study sought to understand how, from the large number of neurotransmitter release sites (active zones, **AZs**), only a very small number of release sites contribute to evoked neurotransmitter release with low variance. We used the toad neuromuscular synapse to address this issue. Using this synapse, we compared the neurotransmitter release characteristics when curare or the Na-channel blocking conotoxin μ-CgTx GIIIB were used to suppress the muscle action potential and contraction as the extracellular calcium concentration [Ca^2+^]_o_ was increased. Under μ-CgTx GIIIB, the maximum transmitter release reached 25 quanta at very high extracellular calcium concentration; conversely, under curare, quantal content reached 148 quanta (*P*=0.0067). Using binomial analysis, we determined that this quantal increase was due to a significant increase in the number of release sites ‘n’ rather than the average probability of release ‘p’. Based on these observations, along with EPC analyses (shape and duration), we propose that a fast autoinhibitory response (< 5 ms from the nerve terminal impulse) limits the maximum level of postsynaptic depolarisation by voltage-dependent regulation of the AChR channel conductance. The following discusses our findings in support of this idea.

### Effect of altering quantal size with curare on the relationship between quantal content and extracellular calcium concentration

In the present study, under curare, *m_i_* had a slope of 15.25 ± 4.75 (R^2^=0.48) when the [Ca^2+^]_o_ was low and increased to 20.11 ± 5.31 (R^2^=0.70) when the [Ca^2+^]_o_ was high (**Figure 1**). One explanation for this increase could be the idea that quantal amplitude may be regulating the release of quanta from the terminal or that quantal amplitude limits the size of the postsynaptic depolarisation. When curare is added, quantal size is reduced and quantal content increased (**Figure 2**). By contrast, when quantal size was not reduced by using µ-CgTx GIIIB, quantal content increased significantly to a lower level when compared to it in the presence of curare. This result further demonstrates that quantal size can greatly influence quantal content.

To examine the possibility that a limiting influence is restricting the level of postsynaptic depolarisation, double reciprocal plots of *m_i_* versus [Ca^2+^]_o_ were constructed to theoretically determine the maximal quantal content when the extracellular calcium approaches infinity (1/[Ca^2+^]_o_ = 0). For curare, quantal content approached 148 compared to 25 in the presence of µ-CgTx GIIIB. The curare result was consistent with previously published data (**Figure 2**, **Table 1**) (Bennett et al., 1977; Dodge & Rahamimoff, 1967). In general, the use of curare to reduce quantal size yielded quantal contents that were closer to the number of AZs in previous published studies and the present study. By contrast, µ-CgTx GIIIB, which does not reduce quantal size, yielded a significantly (*P*=0.0067) lower quantal content. These results demonstrated that quantal size does influence quantal content. Specifically, it supports the idea that there is an inverse relationship between quantal size and quantal content, indicative of potential compensation (Edwards, 2007).

### Effect of curare on quantal content

Recently, the ability of curare and other blockers of postsynaptic AChRs to increase quantal content has been suggested to be a form of presynaptic homeostatic plasticity (Wang et al., 2018; Wang et al., 2016). Namely, in these studies, researchers have proposed that a mechanism operates to upregulate presynaptic neural excitability to compensate for a loss of function (blockage of AChRs) (Wang et al., 2018; Wang et al., 2016). Specifically, Wang and colleagues used binomial analysis to determine that a block of AChRs by α−bungarotoxin results in an upregulation of the binomial parameter ‘n’. In agreement with these findings and proposals, our data, presented in Figure 3, indicates that under both curare and µ-CgTx GIIIB, the probability of release ‘p’ rapidly increases to a maximum at 0.7mM extracellular calcium (**Figure 3B**), and that the further increase in quantal content originates from an increase in binomial parameter ‘n’ (**Figure 3C**).

However, what the physical correlate of binomial parameter ‘n’ is, regarding its exact mechanistic identity, has long been controversial. Several studies have suggested that ‘n’ is best described as a single AZ, while other researchers have proposed that ‘n’ represents a single release-ready synaptic vesicle (Stevens, 2003; Wang et al., 2010). Crucially, an increase in the number of participating synaptic vesicles viewed strictly would suggest a purely presynaptic origin. Conversely, at the neuromuscular synapse, an increase in ‘n’ may arise because of a reduction in the proportion of postsynaptic AChRs opposite an AZ being activated by the release of a quantum of transmitter. If the level of depolarisation of a quantum of transmitter is reduced, this reduction could allow for some not yet activated AChRs to interact with subsequent quantal release, resulting in an observed increase in binomial parameter ‘n’. This possible contribution is because recordings of evoked and spontaneous are ultimately measurements of the activity of the AChRs participating in transmission (i.e., postsynaptic measurements). In support of this notion, recent studies have further emphasised this tight trans-synaptic connection controlling levels of synaptic transmission (Guzikowski & Kavalali, 2021; Tang et al., 2016). Considering these observations and recent assertions, neuromuscular transmitter release sites could be better described as trans-synaptic AZ complexes, involving an interaction between pre-synaptic release of quanta and post-synaptic activation of AChRs; an idea supported by our findings.

While retrograde signals, including trophic factors and basal lamina molecules, secreted from muscle to the pre-synaptic nerve terminal, remain a well-argued possibility to regulate synaptic transmission by a longer time course, supported by functional and molecular evidence (Chand et al., 2015; Fong et al., 2010; Nishimune et al., 2004; Rogers & Nishimune, 2017; Wang et al., 2016; Zhu et al., 2021). A much faster regulation would involve the character of the AChRs. For example, it was found that functional blockage of AChRs caused an increase in the decay rate of mEPCs (Wang et al., 2018) - a phenomenon also observed in past studies on repetitive stimulation under curare (Harborne et al., 1988). The faster decay time after curare administration was the reason why Wang et al (2018) suggested a postsynaptic conformational change of AChRs as a mechanism for their proposed trans-synaptic signalling (Wang et al., 2018). Considering that AChRs with less conductance or shorter opening time (faster decay rate) would cause a smaller influx of cations (Na+) from the extracellular space, supports a role for AChRs in the apparent limitation of quantal release.

### Possible mechanism of postsynaptic autoregulation of quantal content

Instead of triggering upregulation of synaptic transmission following a reduction in quantal size by drugs that partly inhibit the function of AChRs, the proportion of AChRs that are not activated during the duration of the first quanta released may remain receptive to subsequent quanta released and thereby contribute to subsequent postsynaptic responses. In this way, a fast and localized auto-inhibition would be imposed by the characteristics of the AChRs. As outlined in the introduction, a recent conceptual model describes the ability of highly active release sites to impose a surrounding autoinhibition, rendering surrounding sites relatively silent (Ge et al., 2020; Bennett & Robinson, 1990). This effectively renders the surrounding trans-synaptic units to become refractory following the activation of one unit within the group of trans-synaptic units. In this way, active trans-synaptic units are autoregulated, reducing the overall number of active sites along nerve terminal branches and, in effect, raising the overall probability of each trans-synaptic unit.

The reduction in depolarisation induced by curare may disrupt this spatial distribution of trans-synaptic units, in effect reducing the size of each unit and, as a result, increasing the number of trans-synaptic units along terminal branches (**Figure 1**). This proposal would explain the observed increase in the number of active release sites when curare was added, and at the same time could explain the probability of release reaching unity as we have previously shown (**Figure 3A**). Hence, we propose that it remains possible that the AChR component of the trans-synaptic unit has a role in the local fast auto-inhibition. Rapid (1-2 ms) desensitisation of AChRs induced by local depolarisation resulting from the release of the transmitter at one of the release sites within the trans-synaptic unit. Rapidly desensitised AChRs by local (quantal) release of ACh would allow for fast synaptic transmission between nerve terminals and muscle. Desensitised receptors possess a higher affinity for ACh than active receptors (Vyskocil et al., 2009), and thus a significant proportion of further releases of ACh (additional quanta) is blocked by previous ACh molecules or “trapped” at desensitised receptors (Sine & Steinbach, 1984; Vyskocil et al., 2009). This mechanism reduces the probability of repetitive binding upon active receptors, resulting in decreased duration of excitability. Meanwhile, the decreased conductance of AChRs desensitised by repetitive exposure to ACh would also result in a reduction in the EPP amplitude and decay time, decreasing neuronal excitability, “depolarising block” (Gage, 1976). We propose that this effect would be more pronounced at highly active release sites, or at very high extracellular calcium concentrations, where multiple EPC events occur and are capable of saturating surrounding AChRs, “enhancing depolarizing block”.

### The effect of an initial end plate current on subsequent end plate currents that occur within 5 ms: time course and amplitude changes

We examined whether an initial EPC generated by a quantum released influenced the time course and/or amplitude of a subsequent EPC if it occurred within a few milliseconds. The premise is that pre-conditioning of AChRs by the release of a quanta of ACh alters the characteristics of the AChRs, resulting in shortening the duration and the amplitude of the EPC. Highly active release sites might impose a refractory influence around the neighbouring release sites. The magnitude of this refractory influence may offer an explanation relating to the non-uniformity in registering EPCs along amphibian motor nerve terminal branches of quantal transmitter release (Bennett & Lavidis, 1979).

In our toad NMJ study, we found that at highly active sites, when twin EPCs (**Figure 4A**) occur, the second of the pair had both a lower conductance by 15.85% overall (*P*=0.00149), and a lower amplitude by 19.25% (**Figure 4B, C**). Further, this effect seems to persist in cases where more than two EPCs are released, as seen in **Figure 5A**, with a "sawtooth" like formation where subsequent pulses were on average 28.06% ± 0.28 smaller than the first EPC. The presence of a preceding EPC seems to inhibit subsequent EPCs at these highly active sites. Additionally, there appears to be a temporal aspect as well, as seen in **Figure 4B** and **C**, when the interval between the paired EPCs was less than 5ms, the second EPC always had a shorter conductance than the first. Similarly, for multiple releases of quanta (**Figure 5B**), the initial EPC reduced the amplitude of subsequent release events, which was most pronounced at quantal events closest to the initial EPC.

Our observations suggest that at the toad neuromuscular synapse, post-synaptic AChRs have a significant role in regulating the level of depolarisation imposed on a muscle fibre during stimulation by the nerve terminal impulse. Even though a typical motor nerve terminal is composed of hundreds of release sites (active zones) that act as release points for quanta of transmitter, they do not act independently. They act in small groups that are formed by very active release sites, which generate trans-synaptic groups of active zones. This action results in reducing the number (hundreds) of functional release sites to act as just a few (tens) of functional release sites. Such an arrangement may allow for reliable fast synaptic transmission and prevent over-excitation of the AChRs, thus reducing a depolarising block and paralysis of the muscle. Altogether, it appears that at highly active sites, there is a temporal factor that limits subsequent release. The effect of a first EPC upon subsequent conductance suggests a post-synaptic involvement. It appears that when AChRs are activated by released ACh, the AChR channels shorten the period they remain open. It is possible that even when vesicles are released from the AZ, some of these quanta may not register EPCs, since the AChRs may be in a refractory period due to previous activation from quanta released. Such a mechanism may explain the idea of missing quanta and offers a possible mechanism proposed by the fast auto-inhibition theory (Dengyun Ge et al., 2020).

In summary, curare significantly increases the maximum possible quanta release from 25 quanta to 148 quanta. Curare partially inhibits postsynaptic AChRs, decreasing the depolarisation induced by single quanta, and thus decreasing the sphere of influence imposed by the transmitter and subsequent depolarisation. This, in effect, then increases the number of functional trans-synaptic units made up of the pre- and postsynaptic structures. A visual representation of this can be seen in **Figure 7**.

**Figure 6.**
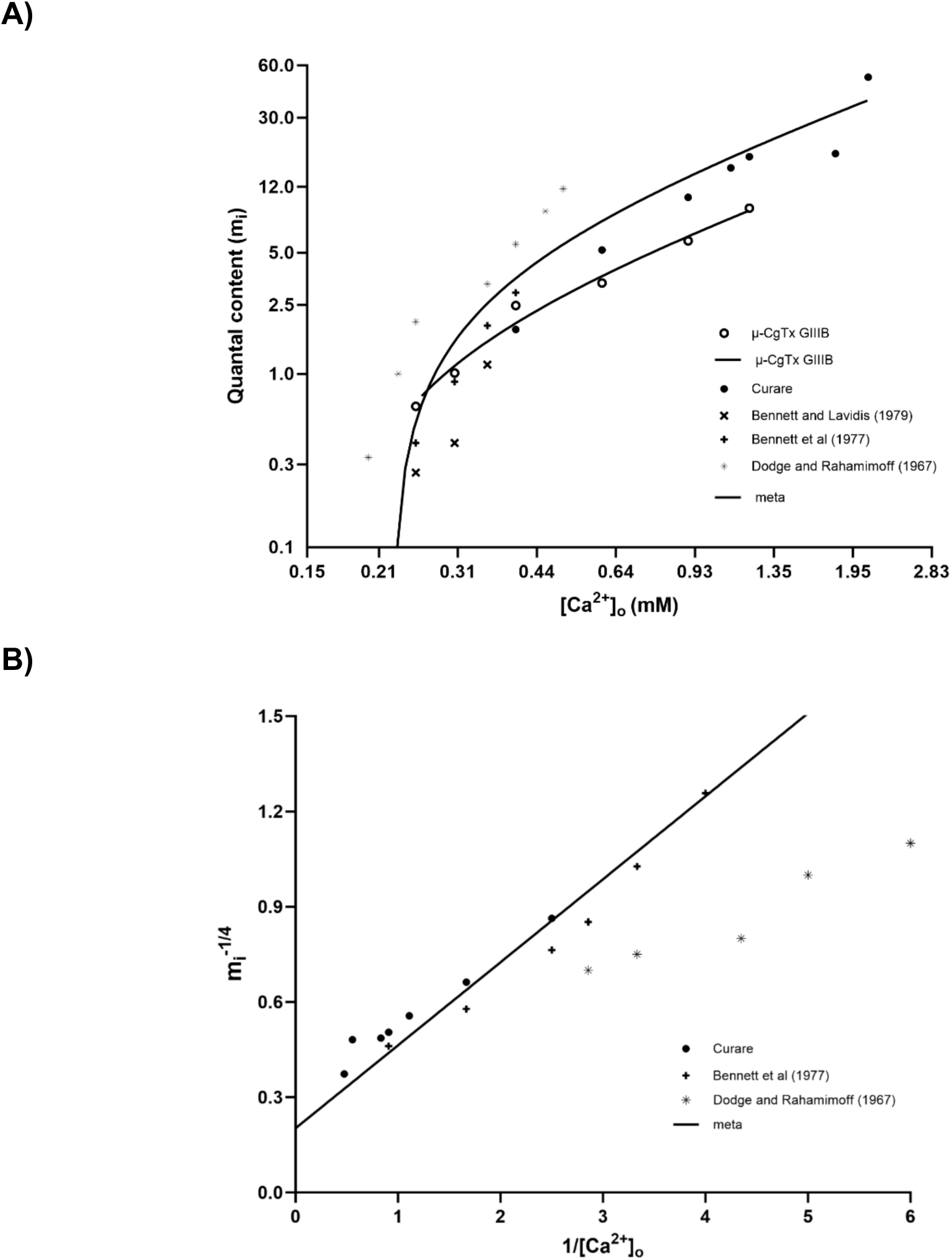
Meta-analysis of curare experiments in the literature. **A)** Meta-analysis of calcium dependence on *m_i_*, and present investigation µ-CgTx GIIIB and curare treatments. A linear regression was constructed from published data, which allowed for comparison to the linear regression for µ-CgTx GIIIB treatment in this study. **B)** Lineweaver-Burk plots constructed using data from the studies shown in Table 1, compared to present results under µ-CgTx GIIIB and curare treatments. At ∞ [Ca^2+^]_o_ linear regression revealed a y-intercept of 0.3045 for Dodge and Rahamimoff (1967), and 0.1709 for Bennett et al (1977). Significant differences between treatments and multiple comparisons determined by Tukey analysis are indicated by asterisks (*P* ≤0.01). Overall, there was an insignificant difference between our results using curare and literature values (*P*=0.8424).

**Figure 7.**
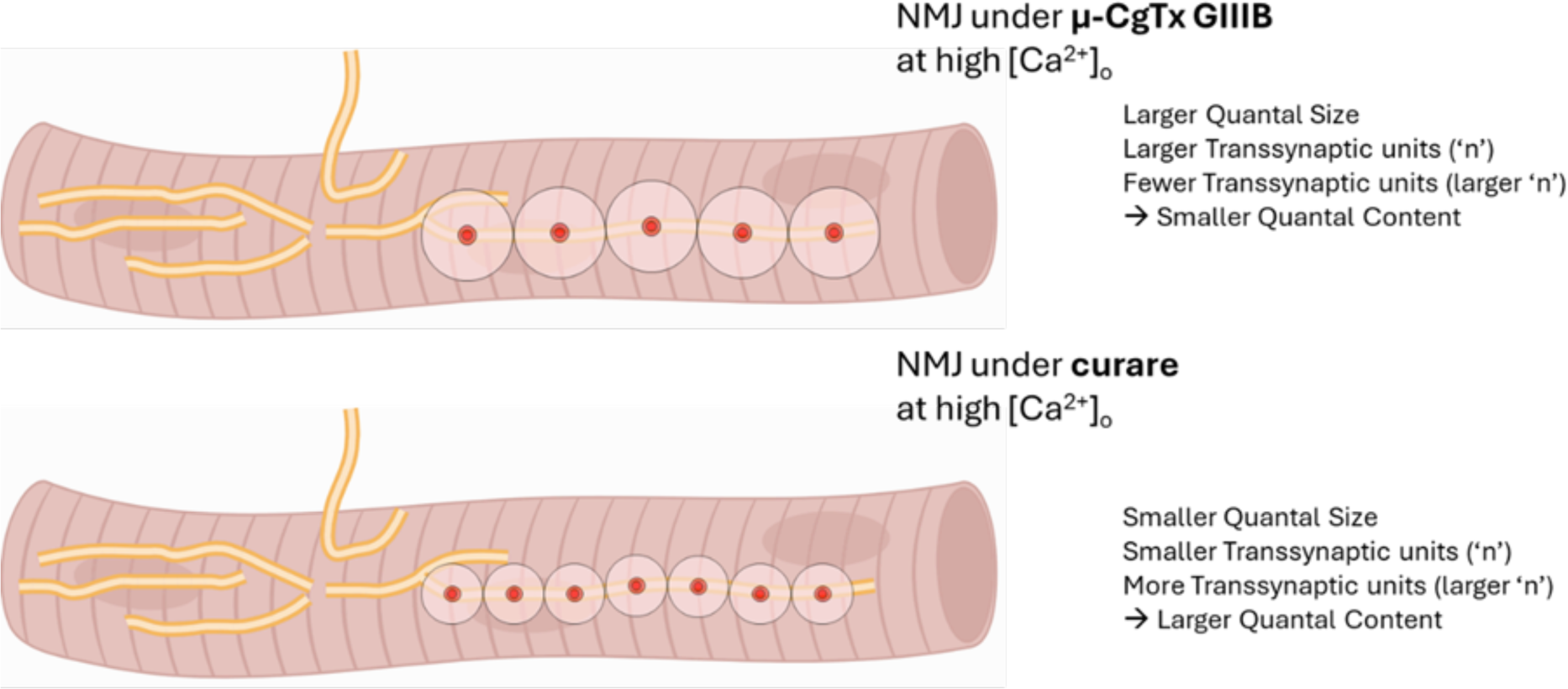
A conceptual diagram describing the relationship between quantal size and quantal content. A reduction in postsynaptic depolarisation, e.g. by curare, disrupts the regular spatial distribution of transsynaptic units (large red and pink circles, top) along the amphibian nerve terminal (yellow structures), reducing the size of each unit (small red and pink circles, bottom), and as a result, increasing the number of transsynaptic units along a terminal branch. Each trans-synaptic unit consists of a group of active zones (red), and their AChRs (dark pink). Inhibitory envelope (pink) adapted from Ge et al., 2020’s fast autoinhibition model.

## Acknowledgements

An Australian Government Research Training Program Scholarship supported this research. Reagents needed for electrophysiology were from PhD support funds and UQ SBMS funds awarded to NAL and PGN.

